# Isolation, identification, and comparative genomic analysis of *Vibrio harveyi* as a causative agent for a skin ulcer disease of endangered fish Chinese bahaba (*Bahaba taipingensis*)

**DOI:** 10.64898/2026.09.10.750546

**Authors:** Liang Zhong, Ruoxi Zhu, Yuxuan Zhang, Lin Yan, Song Sun, Kuoqiu Yan, Wenlong Cai

## Abstract

Chinese bahaba (*Bahaba taipingensis*) is a fish species endemic to China and listed as a Class I nationally protected wild animal. Although significant breakthroughs have been achieved in its artificial breeding, high mortality rates persist during cultivation. Beginning in August 2024, sustained mass mortalities occurred in artificially farmed Chinese bahaba at a farm in Guangdong Province. Diseased fish exhibited pronounced skin ulcers, ascites, gastroenteritis, and splenomegaly. Histopathological examination revealed severe necrotizing gastroenteritis, gill necrosis and edema, muscle congestion with inflammatory cell infiltration, hepatic congestion with marked vasculitis, renal interstitial necrosis with tubular atrophy, and abundant melanomacrophage centers in the spleen. Based on biochemical characterization and whole-genome phylogenomic analysis, the isolated strain HCY1 was identified as *Vibrio harveyi*. Artificial infection experiments with this strain resulted in 100% mortality and reproduced clinical signs similar to those in naturally infected fish. Antimicrobial susceptibility testing showed that HCY1 remained susceptible to florfenicol and trimethoprim-sulfamethoxazole, but was resistant to multiple antibiotics, including β-lactams and tetracyclines. Comparative genomic analysis revealed that HCY1 carries various antimicrobial resistance genes primarily conferring resistance to β-lactam and tetracycline antibiotics, consistent with the phenotypic resistance profile. The strain also harbored numerous virulence genes, including complete type III and type VI secretion systems (T3SS and T6SS), *tlh*, *hap*/*vvp*, and *rtxB*. Its plasmids encoded multiple virulence-enhancing genes, such as *mcpA*, *pal*, and *esiB*. To our knowledge, this is the first systematic investigation of a disease outbreak in cultured Chinese bahaba, and provides valuable insights for disease control and population conservation of this endangered species.

## 1 Introduction

Chinese bahaba (*Bahaba taipingensis*) belongs to the genus *Bahaba*, family Sciaenidae, order Perciformes, and is a fish species endemic to China. It is one of the largest species within the family Sciaenidae, reaching a maximum body length of over 2 meters and a weight of up to 100 kg (Chu et al., 1963). This species is primarily distributed in estuarine waters of China, including the Pearl River Estuary in Guangdong Province, the Hangzhou Bay and Zhoushan waters in Zhejiang Province, and the Minjiang River Estuary and Quanzhou Bay in Fujian Province (Sadovy and Cheung, 2003). This fish species has great ecological, economic, and cultural value. As the largest species in the family Sciaenidae, it occupies a high trophic level (3.9 ± 0.4 se, FishBase) and plays an important role in maintaining the stability of estuarine ecosystems (Chen et al., 2025; Hu et al., 2025). Economically, the price of its swim bladder is tens of thousands of dollars per kilogram, which is far exceeding the prices of swim bladders of other fish species in the family Sciaenidae (Smith et al., 2023). As a result, its swim bladder is known as “one of the world’s nine most expensive foods or medicines”. The consumption of fish swim bladders has a long history in China, especially in southern coastal areas. The swim bladder is traditionally believed to possess high nutritional and medicinal value, containing multiple beneficial substances for humans, such as polysaccharides, elastin, and collagen (Cruz-López et al., 2023; Pan et al., 2018). It is often used as a tonic to aid recovery after surgery or childbirth, with the swim bladder of Chinese bahaba being particularly prized. In the Chaoshan area, residents traditionally keep the swim bladder of Chinese bahaba as a family heirloom, intended to guard against postpartum hemorrhage. In addition, the swim bladder has been used as a gift or investment (Xu et al., 2024).

In recent years, wild populations of Chinese bahaba have declined sharply due to climate change, marine pollution, and overfishing driven by the high value of its swim bladder (Zhao et al., 2012). In 2006, the International Union for Conservation of Nature (IUCN) Red List of Threatened Species classified it as Critically Endangered (CR) (Liu, 2020), and in 2021, it was listed as a Class I nationally protected wild animal (National Forestry and Grassland Administration, 2021). To bolster conservation of this endangered species, artificial breeding of Chinese bahaba began in 2021, with the first successful production of offspring achieved in 2022. Over the following years, multiple batches of artificially bred offspring have been produced. However, high mortality rates have been observed under indoor rearing conditions, despite careful management (Cui et al., 2024). Therefore, to improve the survival rate of artificially reared Chinese bahaba, it is particularly important to investigate diseases affecting farmed individuals.

Vibrios are Gram-negative bacteria characterized by short, curved rod-shaped or comma-shaped morphology and belong to the phylum Proteobacteria, class Gammaproteobacteria, order Vibrionales, and family Vibrionaceae (Sampaio et al., 2022). Vibrios are one of the most ubiquitous bacterial groups in nature, commonly found in marine, estuarine, and freshwater environments (Sampaio et al., 2022). They are particularly prevalent in warm waters, especially when temperatures exceed 17°C (Oliver et al., 2012).To date, more than 100 *Vibrio* species have been identified, most of which are nonpathogenic (Baker-Austin et al., 2018). About 12 *Vibrio* species are known to infect humans, such as *Vibrio cholerae*, *V. parahaemolyticus*, and *V. vulnificus* (Baker-Austin et al., 2018; Ceccarelli et al., 2019). However, some *Vibrio* species are harmful to aquatic animals. For instance, *V. harveyi*, *V. alginolyticus*, and *V. parahaemolyticus* are frequently detected in aquaculture systems, where they cause mass mortalities in cultured species, leading to severe economic losses (Amalina et al., 2019; Deng et al., 2020; Mohamad et al., 2019).

*V. harveyi* is a fermentative, rod-shaped bacterium characterized by polar flagella and bioluminescent properties (Zhang et al., 2020). As one of the most significant members of the genus *Vibrio*, *V. harveyi* is also one of the most common pathogens in aquaculture. It can infect not only vertebrates such as fish, but also invertebrates, including shrimp and shellfish (Muthukrishnan et al., 2019; Wei et al., 2019; Yuan et al., 2021). Furthermore, it can infect cultured aquatic animals at various growth stages, from larvae to adults (Dong et al., 2017; Gauger et al., 2006). Infected animals typically exhibit clinical signs such as ulcerative skin lesions and gastroenteritis (Austin and Zhang, 2006). Additionally, pronounced pathological damage is often observed in multiple tissues and organs, including the muscle, brain, and kidney (Dong et al., 2017), ultimately leading to mass mortality with cumulative mortality rates reaching as high as over 80% (Soto-Rodriguez et al., 2012). This indicates that *V. harveyi* poses a great threat to aquaculture.

In this study, we reported a mass mortality event in artificially cultured Chinese bahaba infected with *V. harveyi* at a farm in the coastal area of Guangdong Province. Through bacteriological, histopathological, and comparative genomic analyses, we systematically elucidate the etiology of this disease outbreak for the first time and provide valuable insights for disease control and conservation of this critically endangered species.

## 2 Materials and Methods

### 2.1 Necropsy

From October 2024 to January 2025, a continuous mortality event occurred among one-year-old artificially bred Chinese bahaba (*Bahaba taipingensis*) (weight: 37-165 g, length: 17-27 cm) reared in a recirculating aquaculture system along the coast of Guangdong Province (License for artificial breeding of aquatic wild animals (*B. taipingensis*) of PRC #2024YL0028), with a cumulative mortality exceeding 20%. Fish in almost all tanks suffered mortality and displayed the same clinical signs. The mortality rate reached its maximum in late October, with daily mortalities reaching hundreds of fish at the peak. During this period, the fish were fed manually three times daily under the following water conditions: dissolved oxygen > 5 mg/L, temperature 24.0-28.7□, salinity 30-33 ppt, ammonia nitrogen < 0.1 mg/L, nitrite < 0.1 mg/L, and pH 7.2-8.0. Necropsy examinations were conducted on freshly dead specimens with typical clinical signs from different tanks. Skin and gill samples from each fish were examined for parasites by microscope, and liver, spleen, and kidney tissues were tested for common viral pathogens by PCR (data not shown), including iridovirus and nervous necrosis virus. No parasites and viruses were observed and tested.

### 2.2 Bacterial isolation

To isolate bacteria, tissue samples were collected from the liver, spleen, kidney, and muscles of the diseased fish. These samples were then streaked onto brain-heart infusion (BHI) agar plates and incubated at 28°C for 24 hours. The isolates were sub-cultured under the same conditions to ensure purity. Following purification, the isolates were suspended in 20–25% glycerol and stored at –80°C.

### 2.3 Bacterial identification

The bacteria were revived by culturing in BHI broth at 28°C for 24 hours. Genomic DNA was then extracted using a Genomic DNA Extraction Kit (Servicebio, Wuhan, China). For species identification, universal 16S rRNA primers (27F: 5’-AGAGTTTGATCCTGGCTCAG-3′ and 1492R: 5’-GGTTACCTTGTTACGACTT-3′) were employed. The PCR protocol consisted of an initial denaturation at 98°C for 2 minutes, followed by 30 cycles of 98°C for 20 seconds, 55°C for 20 seconds, and 72°C for 10 seconds, with a final extension step at 72°C for 5 minutes. The amplified sequences were compared using BLAST against the GenBank database (https://www.ncbi.nlm.nih.gov/) for identification of the bacterial isolates.

### 2.4 Biochemical characterization

The purified bacterial strain was revived on Tryptic Soy Agar (TSA) and subsequently used for biochemical characterization. Biochemical properties of the strain were assessed using the API 20NE test kit (bioMérieux, France), following the manufacturer’s instructions. The tested biochemical characteristics included potassium nitrate reduction, glucose fermentation, arginine dihydrolase, urea hydrolysis, esculin/aesculin hydrolysis, gelatin hydrolysis, β-galactosidase activity (o-Nitrophenyl-β-D-galactopyranoside/o-NPG), assimilation of glucose, L-Arabinose, mannose, mannitol, N-Acetylglucosamine (GlcNAc), maltose, gluconate, capric acid (decanoic acid), adipic acid, malic acid, citric acid, phenylacetic acid, and tryptophan-based indole production.

### 2.5 Antibiotic susceptibility tests

The disc diffusion method was employed to assess the antibiotic susceptibility of the isolated strain. A total of twenty kinds of antibiotics were tested, including Penicillin (10U/disc), Ampicillin (10 μg/disc), Oxacillin (1 μg/disc), Ceftriaxone (30 μg/disc), Cefalexin (30 μg/disc), Kanamycin (30 μg/disc), Gentamycin (10 μg/disc), Amikacin (30 μg/disc), Neomycin (30 μg/disc), Tetracycline (30 μg/disc), Doxycycline (30 μg/disc), Erythromycin (15 μg/disc), Minocycline (30 μg/disc), Ofloxacin (5 μg/disc), Enrofloxacin (10 μg/disc), Florfenicol (30 μg/disc), Chloramphenicol (30 μg/disc), Clindamycin (2 μg/disc), Vancomycin (30 μg/disc), and trimethoprim-sulfamethoxazole (23.75/1.25 μg/disc). In brief, the bacterial isolate was collected, resuspended in sterile phosphate-buffered saline (PBS), and adjusted to a concentration of 1 × 10 CFU/mL. The obtained bacterial suspension was then spread onto BHI agar plates. Following incubation at 28°C for 24 hours, the diameters of the inhibition zones were measured. The isolates were categorized as resistant, intermediate, or susceptible based on the criteria outlined in the Clinical and Laboratory Standards Institute (CLSI, 2018).

### 2.6 Histopathological analysis

Various tissues from the diseased fish (liver, spleen, kidney, head kidney, heart, stomach, intestine, brain, muscle, and swim bladder) were immersed in 10% neutral buffered formalin for 48 hours to achieve fixation. Afterwards, the specimens were placed into cassettes, rinsed under running water overnight, and then subjected to dehydration through a graded ethanol series, clearing in xylene, and infiltration with paraffin wax. Tissue sections of 4 μm thickness were cut and stained with hematoxylin and eosin (H&E) for subsequent microscopic examination (Liu et al., 2023).

### 2.7 Fish challenge

Due to the endangered status of Chinese bahaba, live individuals could not be used as subjects for challenge experiments. Therefore, we selected its closest phylogenetic relative, the Miiuy croaker (*Miichthys miiuy*), as a surrogate species for artificial infection (Cui et al., 2024). Healthy Miiuy croaker (body weight: 25.33 ± 6.12 g, body length: 10.52 ± 0.82 cm) were obtained from a farm in Guangdong Province. Before the challenge experiments, the fish underwent a two-week acclimation period to adjust to the new environment. To confirm the absence of pre-existing bacterial infection, three fish were randomly selected, and bacterial isolation was attempted from their liver, spleen, and kidney. Only fish that yielded no bacterial isolates were deemed suitable for the subsequent challenge trials. During the acclimation period, continuous aeration was provided to maintain dissolved oxygen levels above 5 mg/L, along with a pH range of 7–8 and a water temperature of 27–28°C. The fish were fed commercial feed twice daily, at 9:00 am and 6:00 pm. A total of 90 fish were randomly allocated into two groups (control and experimental groups), each consisting of three replicates with 15 fish per replicate.

Before the artificial challenge, the fish were fasted for one day. The isolated strain HCY1 was grown overnight in BHI broth at 28°C, then harvested by centrifugation at 4000 rpm for 10 minutes and resuspended in sterile PBS. Fish in the experimental group received an intraperitoneal injection of 0.1 mL of this bacterial suspension with a final concentration of 2.3×10^6^ CFU/g according to our LD_50_ assays, whereas those in the control group were injected with the same volume of sterile PBS. Daily observations were made to record clinical signs and mortality. Bacterial re-isolation was performed from the liver, spleen, and kidney of moribund fish. All procedures involving animals were approved by the Animal Care and Use Committee of the City University of Hong Kong, complying with the university’s guidelines for animal experiments.

### 2.8 DNA extraction, sequencing, and De novo genome assembly

After growing strain HCY1 in BHI broth for 24 hours, its genomic DNA was extracted using the TaKaRa MiniBEST Bacteria Genomic DNA Extraction Kit Ver. 3.0 (Takara, Japan) according to the manufacturer’s standard protocol. The quantity, purity, and integrity of the extracted DNA were evaluated using a NanoDrop Spectrophotometer (Thermo Fisher Scientific) and 1.5% agarose gel electrophoresis. From 1 µg of high-quality genomic DNA, Nanopore libraries were prepared with the Ligation Sequencing gDNA-Native Barcoding Kit 24 V14 (SQK-NBD-114.24, ONT, Oxford, United Kingdom). Long-read sequencing was carried out on a MinION (ONT) platform equipped with an R.10.4.1 Flow Cell (FLO-MIN114, ONT). The resulting pod5 files underwent base-calling using Dorado v0.4.1 (https://github.com/nanoporetech/dorado) with the “dna_r10.4.1_e8.2_400bps_sup@v4.2.0” model (Gao et al., 2026).

Statistics on the long-read sequencing data were generated using NanoPlot v1.41.6. Before assembly, the ONT reads obtained from simplex basecalling were filtered with Filtlong, applying the parameters -min-mean-q 80 and -min_length 1000. The obtained high-quality reads were then assembled using Flye v2.9.4 under default settings (Zhang and Austin, 2005; Zhang et al., 2025). The assembled genome was subsequently polished with long reads using Medaka v1.11.3 (https://github.com/nanoporetech/medaka). Assessment of the completeness of the polished assemblies was performed with BUSCO (Benchmarking Universal Single-Copy Orthologs) v5.7.1 (Manni et al., 2021). Functional annotation of the genome assembly was carried out using Prokka v1.14.6 (Seemann, 2014), with gene prediction conducted by Prodigal v2.6.3 (Hyatt et al., 2010). The complete genome sequence of strain HCY1 has been deposited in the National Center of Biotechnology Information (NCBI) database under accession number PRJNA1356466.

### 2.9 The whole genome phylogenetic tree

A whole-genome phylogenetic tree was constructed using 19 strains in total, comprising 6 isolates of *V. harveyi* and 13 other common *Vibrio* species (*V. harveyi* SB1 (GCF_030060435.1), *V. harveyi* 45T2 (GCF_039542195.1), *V. harveyi* N8T11 (GCF_039544385.1), *V. harveyi* ATCC 33843 (GCF_000770115.1), *V. alginolyticus* E110 (GCF_023650915.1), *V. parahaemolyticus* JD2305 (GCF_050920865.1), *V. vulnificus* ATCC 27562 (GCF_002224265.1), *V. cholerae* RFB16 (GCF_008369605.1), *V. campbellii* BoB-53 (GCF_002906475.1), *V. rotiferianus* B64D1 (GCF_002214395.1), *V. natriegens* ATCC 14048 (GCF_035621455.1), *V. azureus* LC2-005 (GCF_002849855.1), *V. mytili* NH-13 (GCF_049563345.1), *V. owensii* DX190301 (GCF_049816615.1), *V. jasicida* 090810c (GCF_002887615.1), *V. diabolicus* ZF102 (GCF_054111865.1), *V. sagamiensis* NBRC 104589 (GCF_007990935.1), in addition to our two isolates *V. harveyi* ZLBH1 and HCY1. Strain ZLBH1 was isolated in our laboratory from Chu’s croaker (*Nibea coibor*) and exhibits high virulence and multidrug resistance (Zhong et al., 2026). All other complete genome sequences were retrieved from the NCBI database. To ensure consistent annotation, all genome sequences were processed with Prokka v1.14.6 with default parameters (Seemann, 2014). Roary (Page et al., 2015) was used to predict orthologous clusters and core genes, with a 95% minimum amino acid identity for clustering and core genes defined as sequences conserved in ≥ 99% isolates. Single-copy core gene families were aligned using MAFFT integrated in the Roary pipeline to generate a concatenated core-genome matrix. Finally, a maximum-likelihood phylogenetic tree was constructed using IQ-TREE 2 (v2.2.2.7) under the LG+G+I substitution model (Nguyen et al., 2014). Node confidence and topology reliability were assessed using 1,000 replicates of the ultra-fast bootstrap (UFBoot) algorithm. The resulting rooted phylogenetic tree was visualized and annotated using iTOL (https://itol.embl.de).

### 2.10 Comparative genomic analysis

For whole-genome multiple alignment and visualization of the three strains, Mauve was used (Darling et al., 2004). Circular genome maps were generated using Proksee (Grant et al., 2023). Antimicrobial resistance (AMR) and virulence genes were analyzed by employing the BLAST tool against the Comprehensive Antibiotic Resistance Database (CARD) and the Virulence Factor Database (VFDB), respectively.

## 3 Results

### 3.1 Necropsy findings

Chinese bahaba of varying sizes occurred significant mortality, with a cumulative mortality rate exceeding 20%. The diseased fish exhibited distinct ulcers on the body surface (Fig. 1A). At the ulcerated sites, scales were extensively detached, leaving the skin completely exposed, with evident congestion and hemorrhage (Fig. 1B). Additionally, the caudal peduncle showed noticeable scale loss along with congestion and hemorrhage (Fig. 1C). The internal organs also exhibited pathological changes of varying extents. A substantial amount of yellow ascitic fluid in the abdominal cavity was observed (Fig. 1D). The liver displayed petechial hemorrhages, while the spleen appeared blunt and enlarged (Fig. 1E). The intestines were swollen with thinned walls and filled with yellow purulent fluid (Fig. 1F).

**Figure 1:**
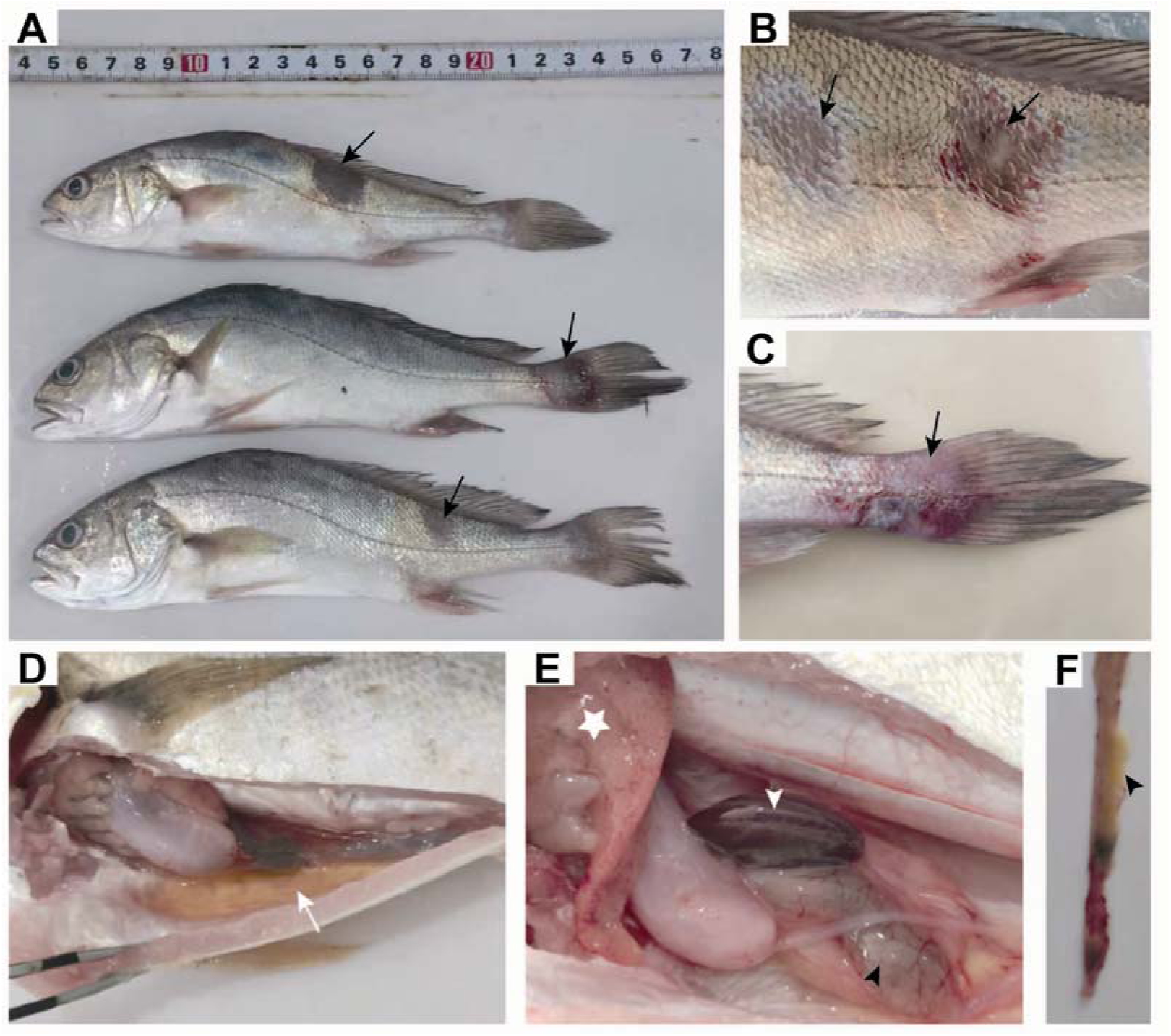
Clinical signs observed in diseased Chinese bahaba. A: Dead fish of various sizes all displayed obvious skin ulcers on the body surface (bland arrow). B: Scale loss, exposed skin, accompanied by significant congestion and hemorrhage, forming ulcerative lesions (black arrow). C: Ulceration on the caudal peduncle, with scale loss, congestion, and hemorrhage (black arrow). D: A large amount of yellow ascitic fluid in the abdominal cavity (white arrow). E: Petechial hemorrhages on the liver (pentagram), and an enlarged spleen with blunt edges (white arrowhead), swollen intestines (black arrowhead). F: The intestinal lumen was filled with pale yellow purulent fluid (black arrowhead).

### 3.2 Histopathology

To further assess the pathological damage in various tissues and organs of the diseased fish, histopathological examination was conducted. The results indicated that the gastrointestinal tract was the primary target organ in the affected fish. In the stomach, the mucosal organization structure was completely disrupted, with severe necrosis of the mucosal layer. Gastric glands showed atrophy, necrosis, and disappearance (Fig. 2A). A large number of necrotic cells had sloughed off and accumulated within the gastric lumen, accompanied by nuclear pyknosis and fragmentation, along with significant inflammatory cell infiltration (Fig. 2B). The intestine exhibited similarly severe necrotic enteritis. The intestinal structure was entirely disorganized, with complete necrosis and sloughing of the mucosal layer (Fig. 2C). Numerous necrotic cells were scattered within the intestinal lumen, displaying nuclear fragmentation, and were accompanied by inflammatory cell infiltration (Fig. 2D). The gills exhibited moderate to severe necrosis and edema (Fig. 2E), with the lamellar bases dissociating from the branchial cartilage and epithelial cells of the lamellae sloughing off (Fig. 2F). The loose connective tissue layer in muscle showed loose tissue structure, congestion, and infiltration of inflammatory cells (Fig. 2G and 2H). In the liver, pancreatic tissue displayed vascular congestion (Fig. 2I), and inflammatory cell infiltration into blood vessels led to vasculitis (Fig. 2J). The kidney exhibited mild interstitial cell necrosis with nuclear fragmentation, atrophy of renal tubules, and detachment of tubular epithelial cells from the basement membrane (Fig. 2K). The spleen contained numerous melanomacrophage centers (Fig. 2L).

**Figure 2:**
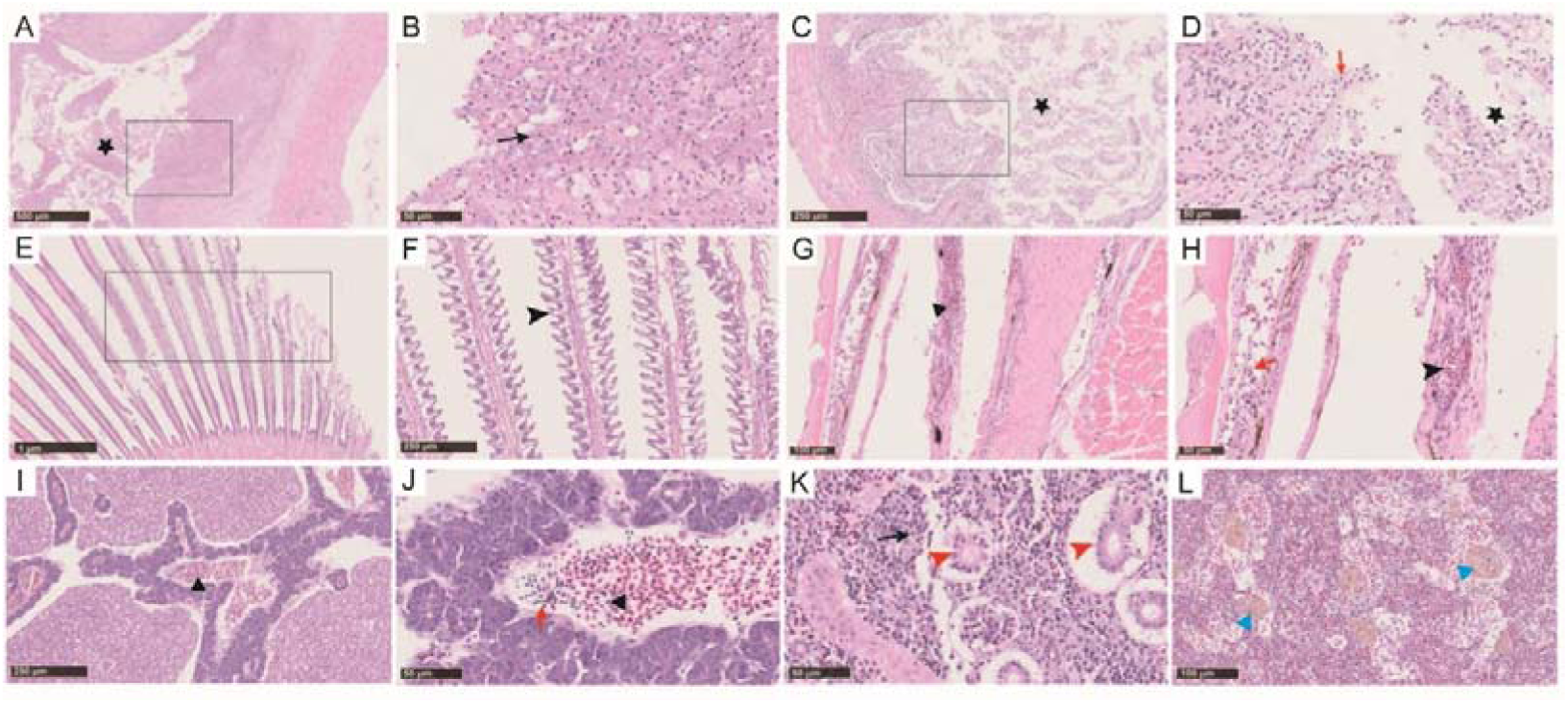
Histopathological observations of diseased Chinese bahaba. A: Severe necrosis and sloughing of gastric mucosa (pentagram), with atrophy and loss of gastric glands. B: Necrotic gastric mucosal layer with pyknotic and fragmented nuclei (black arrow), accompanied by inflammatory cell infiltration. C: Severe necrosis of the intestinal mucosa, disrupted structure, and numerous necrotic cells shed into the intestinal lumen (pentagram). D: Necrotic and detached mucosal cells (pentagram) with inflammatory cell infiltration (red arrow). E: Moderate to severe necrosis in the gill. F: Edema and epithelial lifting of gill lamellae (arrowhead). G: Loose tissue structure and congestion (black triangle) in the loose layer of the dermis of the muscle. H: Muscle congestion (black triangle) and inflammatory cell infiltration (red arrow). I: Pancreatic congestion (black triangle). J: Inflammatory cell infiltration (red arrow) within pancreatic blood vessels (black triangle). K: Necrosis of renal interstitial cells with nuclear fragmentation (black arrow), renal tubular atrophy, and detachment of tubular epithelial cells from the basement membrane (red arrowhead). L: Melanomacrophage centers in the spleen (blue triangle).

### 3.3 Biochemical characteristics

The biochemical characteristics of the strains were analyzed using the API 20NE galleries, and the results were presented in Table 1. The biochemical profile of isolate HCY1 was largely consistent with that of the reference strain CECT 525 that was published by Schrama et al.(2025), except for discrepancies in three phenotypes (urea, L-Arabinose, and citric acid utilization). Both strains were positive for potassium nitrate, glucose, esculin/aesculin, gelatin, O-Nitrophenyl-β-D-galactopyranoside/o-NPG, mannitol, maltose, gluconate, malic acid, while they were negative for arginine, decanoic acid, adipic acid, and phenylacetic acid.

**Table 1:** Results of API 20NE characteristics of strain HCY1.

| Items | HCY1 | CECT 525* |
| --- | --- | --- |
| Potassium nitrate (NO<sub>3</sub>) | + | + |
| Glucose (GLU) | + | + |
| Arginine (ADH) | - | - |
| Urea (URE) | + | - |
| Esculin/aesculin (ESC) | + | + |
| Gelatin (GEL) | + | + |
| O-Nitrophenyl-β-D-galactopyranoside/o-NPG (PNG) | + | + |
| Glucose (GLU) | + | + |
| L-Arabinose (ARA) | + | - |
| Mannose (MNE) | + | +/- |
| Mannitol (MAN) | + | + |
| N-Acetylglucosamine/GlcNAc (NAG) | - | +/- |
| Maltose (MAL) | + | + |
| Gluconate (GNT) | + | + |
| Decanoic acid (CAP) | - | - |
| Adipic acid (ADI) | - | - |
| Malic acid (MLT) | + | + |
| Citric acid (CIT) | - | + |
| Phenylacetic acid (PAC) | - | - |
| Tryptophan (TRP) | + | N |

Note: +, positive; -, negative; ND, not determined. \*: The detection data were sourced from
Schrama et al (2025).

### 3.4 Whole-genome phylogenetic analysis

To clarify the phylogenetic position of the isolated strain HCY1, we constructed a maximum-likelihood phylogenetic tree based on the whole-genome sequence and those of common *Vibrio* species retrieved from the NCBI database. Phylogenetic analysis revealed that the genomic sequence of HCY1 showed high homology with other strains of *V. harveyi* (Fig. 3). These results confirmed that the isolate HCY1 is *V. harveyi*.

**Figure 3:**
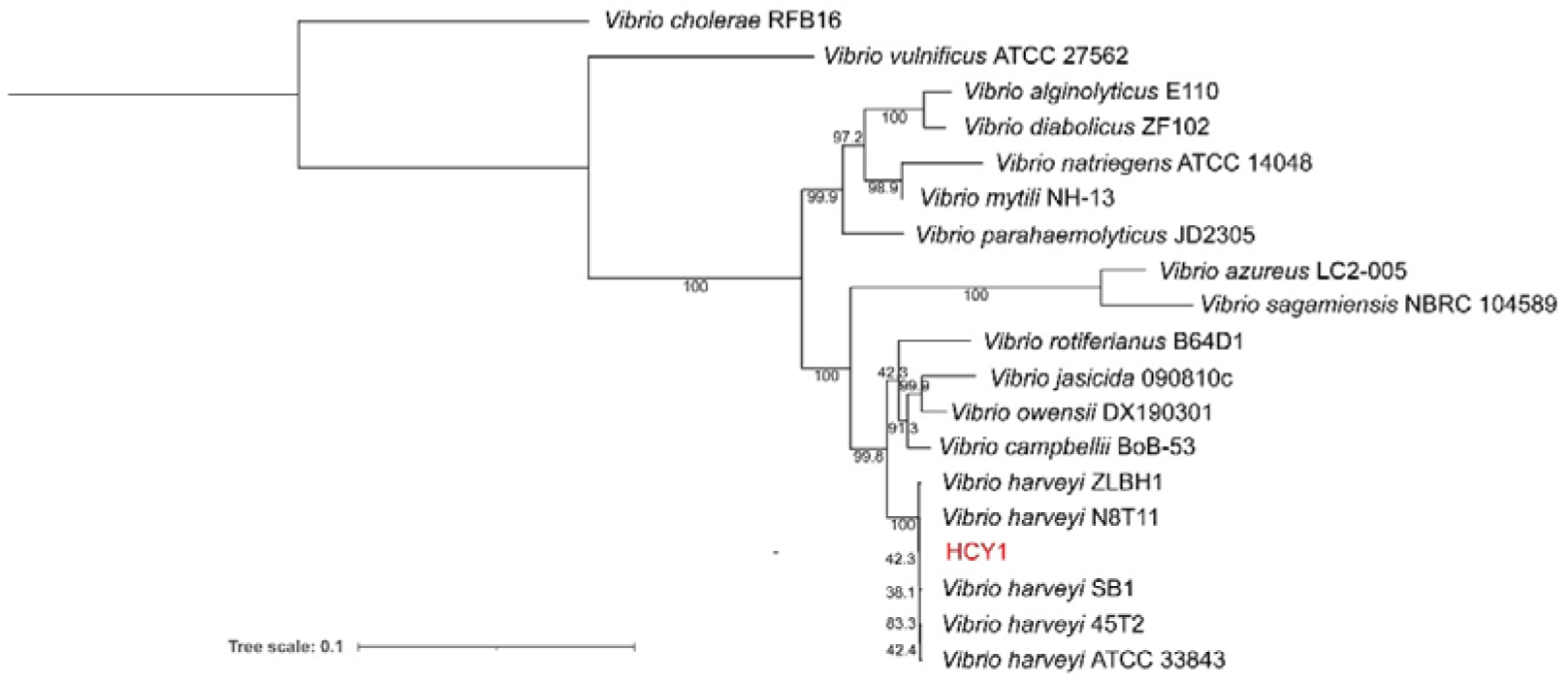
Phylogenetic tree of strain HCY1 using the maximum-likelihood method. Phylogenetic tree based on the whole genome sequences of HCY1 and the other *Vibrio* species. Bootstrap support values were shown at the nodes. *V. cholerae* RFB16 was included as an outgroup to root the phylogenetic tree. The whole genome sequence of HCY1 was deposited in the National Center for Biotechnology Information (NCBI) at https://www.ncbi.nlm.nih.gov/ (Accession numbers: PRJNA1356466). The whole genome sequences of closely related species were obtained from NCBI.

### 3.5 Antibiotic susceptibility tests

To further characterize the antibiotic resistance profile of the isolate HCY1, antimicrobial susceptibility testing was performed (Table 2). The results revealed that the strain remained susceptible to ceftriaxone, gentamycin, amikacin, and chloramphenicol. Furthermore, the strain was susceptible to florfenicol and trimethoprim-sulfamethoxazole, two antimicrobial agents commonly used in aquaculture, suggesting that these may be effective for treating this disease. However, the strain exhibited high-level resistance to β-lactams and tetracyclines. Additionally, resistance was observed against cefalexin, minocycline, enrofloxacin, clindamycin, and vancomycin.

**Table 2:** Antibiotic sensitivity of strain HCY1.

| Antibiotics | Concentration | R/mm | I/mm | S/mm | Inhibition<br>Zone<br>Diameter<br>(mm) | Susceptibility |
| --- | --- | --- | --- | --- | --- | --- |
| **β-lactams** |  |  |  |  |  |  |
| Penicillin | 10U/disc | $\leq 14$ | 15–19 | $\geq 20$ | 0 | R |
| Ampicillin | 10 μg/disc | $\leq 13$ | 14–16 | $\geq 17$ | 0 | R |
| Oxacillin | 1 μg/disc | $\leq 10$ | 11–12 | $\geq 13$ | 0 | R |
| Ceftriaxone | 30 μg/disc | $\leq 13$ | 14–21 | $\geq 22$ | 28 | S |
| Cefalexin | 30 μg/disc | $\leq 14$ | 15–17 | $\geq 18$ | 0 | R |
| **Aminoglycosides** |  |  |  |  |  |  |
| Kanamycin | 30 μg/disc | $\leq 13$ | 14–17 | $\geq 18$ | 16 | I |
| Gentamycin | 10 μg/disc | $\leq 12$ | 13–14 | $\geq 15$ | 15 | S |
| Amikacin | 30 μg/disc | $\leq 14$ | 15–16 | $\geq 17$ | 17 | S |
| Neomycin | 30 μg/disc | $\leq 12$ | 13–16 | $\geq 17$ | 13 | I |
| **Tetracyclines** |  |  |  |  |  |  |
| Tetracycline | 30 μg/disc | $\leq 14$ | 15–18 | $\geq 19$ | 0 | R |
| Doxycycline | 30 μg/disc | $\leq 12$ | 13–15 | $\geq 16$ | 0 | R |
| **Macrolides** |  |  |  |  |  |  |
| Erythromycin | 15 μg/disc | $\leq 13$ | 14–22 | $\geq 23$ | 14 | I |
| Minocycline | 30 μg/disc | $\leq 14$ | 15–18 | $\geq 19$ | 0 | R |
| **Quinolones** |  |  |  |  |  |  |
| Ofloxacin | 5 µg/disc | ≤12 | 13-15 | ≥16 | 14 | I |
| Enrofloxacin | 10 µg/disc | ≤16 | 17-22 | ≥23 | 10 | R |
| Florfenicol | 30 µg/disc | ≤12 | 13-17 | ≥18 | 22 | S |
| Chloramphenicol | 30 µg/disc | ≤12 | 13-17 | ≥18 | 22 | S |
| Clindamycin | 2 µg/disc | ≤14 | 15-20 | ≥21 | 0 | R |
| **Glycopeptide** |  |  |  |  |  |  |
| Vancomycin | 30 µg/disc | ≤14 | 15-16 | ≥17 | 0 | R |
| **Sulfonamides** |  |  |  |  |  |  |
| Trimethoprim-sulfamethoxazole (SMZ/TMP) | 23.75/1.25 | ≤10 | 11-15 | ≥16 | 19 | S |

Notes: S: susceptible, I: intermediate, R: resistant.

### 3.6 Fish challenging

The isolate HCY1 was used to artificially infect the Miiuy croaker. Mortality began within 24 hours post-infection, and the cumulative mortality reached 100% within ten days (Fig. 4A). Moribund fish exhibited clinical signs similar to those observed in naturally infected individuals, including desquamation of scales, exposed skin, ascites, as well as congestion and hemorrhage in the gastrointestinal tract (Fig. 4B-4D). These findings indicate that HCY1 possesses strong virulence.

**Figure 4:**
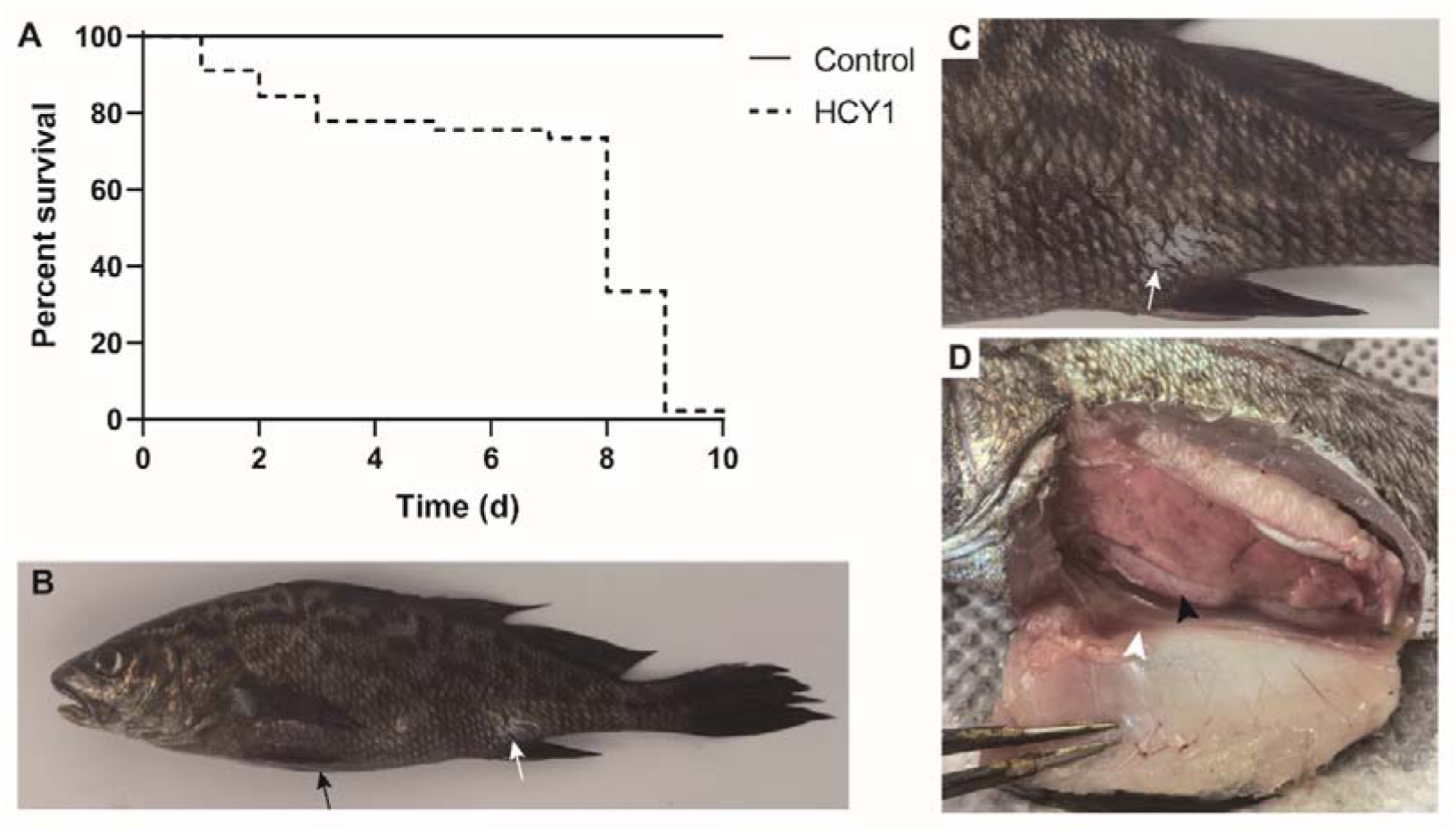
Cumulative mortality and clinical symptoms of artificially infected Miiuy croaker. A: Survival rates of different infection groups. B-D: Artificially infected Miiuy croaker exhibited clinical signs including loss of scales (white arrow), abdominal distension (black arrow), ascites (white arrowhead), and congestion with hemorrhage in the gastrointestinal tract (black arrowhead).

### 3.7 Comparative genomic analysis of *V. harveyi*

To understand the genomic characteristics of this *V. harveyi* HCY1 strain, the complete genome sequence was obtained using Nanopore sequencing. A total of 2,630,297 reads were generated, totaling 1,563,393,716 bp in length, with the longest length reaching 303,284 bp, average length reaching 594.4 bp, a GC content of 44.37%, and a sequencing coverage of 254.02× (Table S1). Following quality control, the complete genome sequence of HCY1 was successfully assembled, comprising two ring chromosomes (3,588,647 bp and 2,250,360 bp) and two plasmids (292,918 bp and 22,673 bp) (Fig. 5).

**Figure 5:**
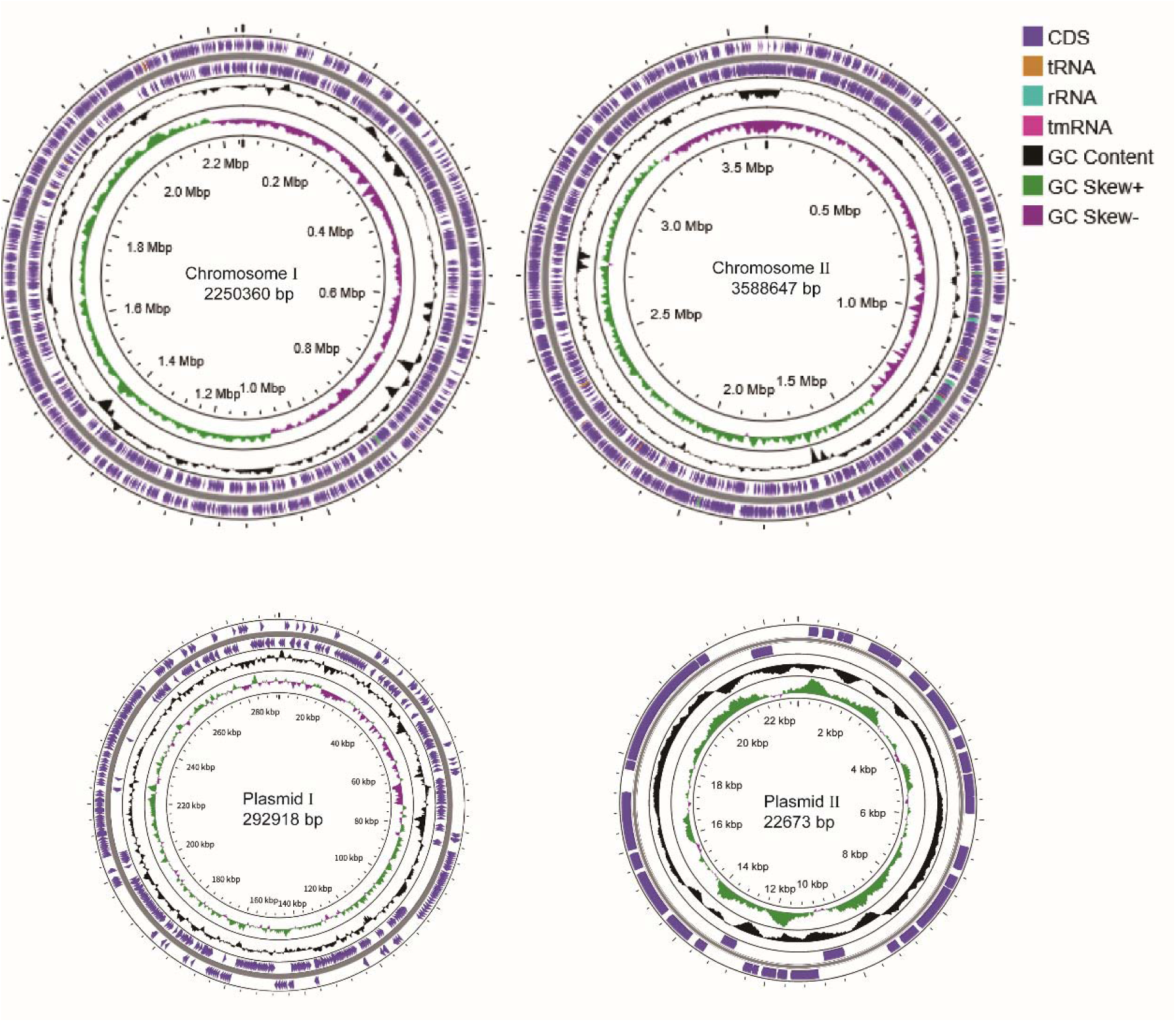
Circular genome of strains HCY1. From the outermost to the innermost ring: the first and third rings represent coding sequences (CDSs), tRNAs, rRNAs, and mRNAs on the two different DNA strands, respectively; the second ring indicates genome size; the fourth ring represents GC content; and the innermost ring displays GC skew.

Two *V. harveyi* strains, including strain 45T2 (low virulence, isolated from large yellow croaker and plasmid-free) and strain ZLBH1 (high virulence, isolated from Chu’s croaker and plasmid-free), were selected for comparative genomic analysis with HCY1. The results showed that the genome of strain HCY1 was highly similar to those of the other two strains, but exhibited obvious translocations and inversions (Fig. 6).

**Figure 6:**
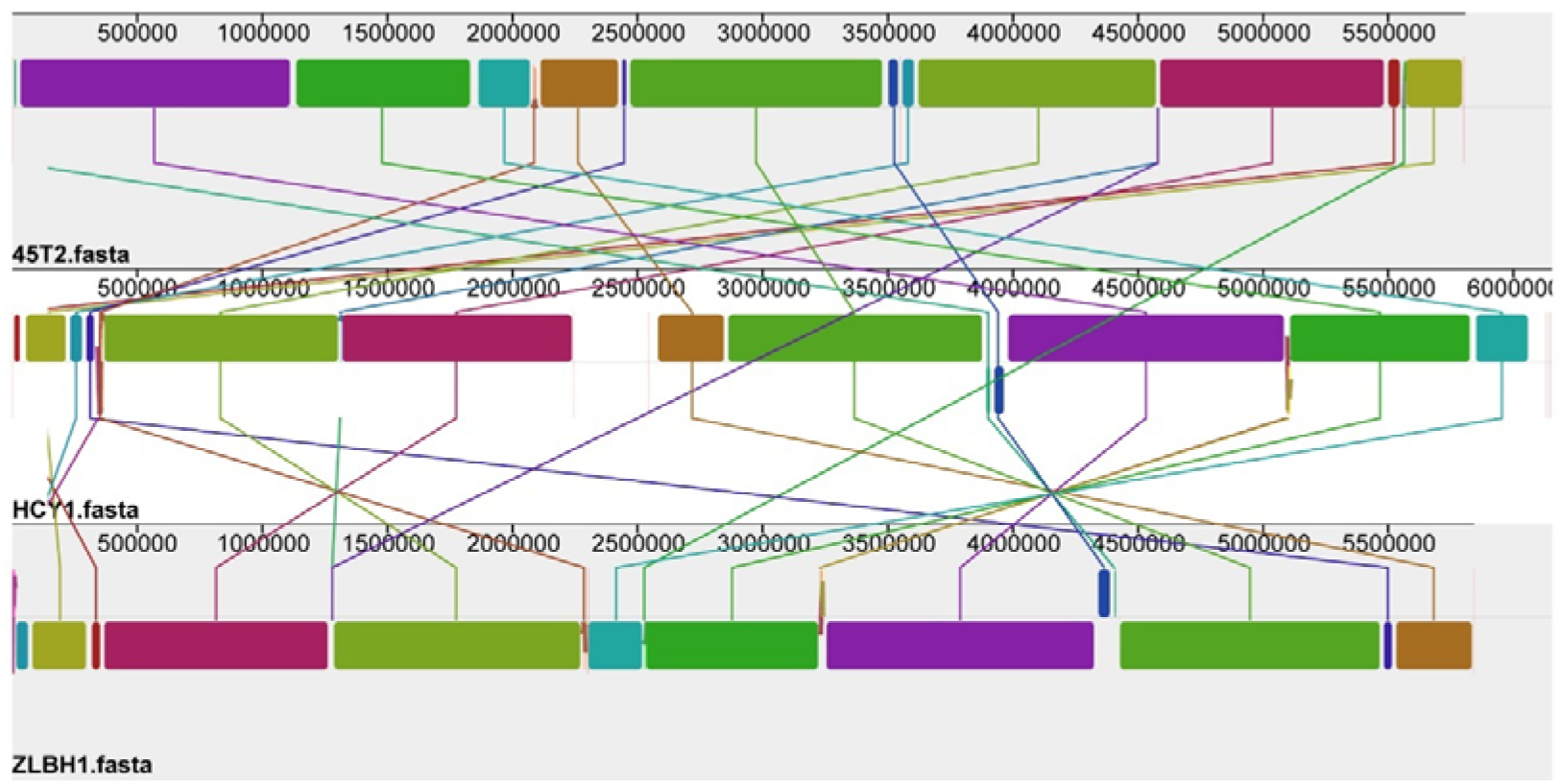
Mauve alignment of 45T2, HCY1, and ZLBH1. Comparison of the linear chromosomal organization among the three *V. harveyi* strains.

### 3.8 Antimicrobial resistance genes analysis

The antimicrobial resistance (AMR) genes were further analyzed (Table 3). The results showed that all AMR genes of HCY1 were located on the chromosomes. And strain HCY1 harbored the highest number of resistance genes (8 AMR genes), with tet(35) and TxR present in two copies each, whereas the low-virulence strain 45T2 carried only five AMR genes, and ZLBH1 carried seven AMR genes. Among these AMR genes, four were shared by all three strains (*tet(35)*, *rsmA*, *CRP*, *VHH-1*). *TxR* was found only in HCY1 and 45T2, while ZLBH1 harbored a unique resistance gene (*ugd*). Notably, a distinct AMR gene, *leuO*, was identified in HCY1, which confers resistance not only to nucleoside antibiotics but also to disinfectants and antiseptics. The AMR genes carried by HCY1 conferred resistance to β-lactams and tetracyclines, which was highly consistent with the antimicrobial susceptibility testing results.

**Table 3:**
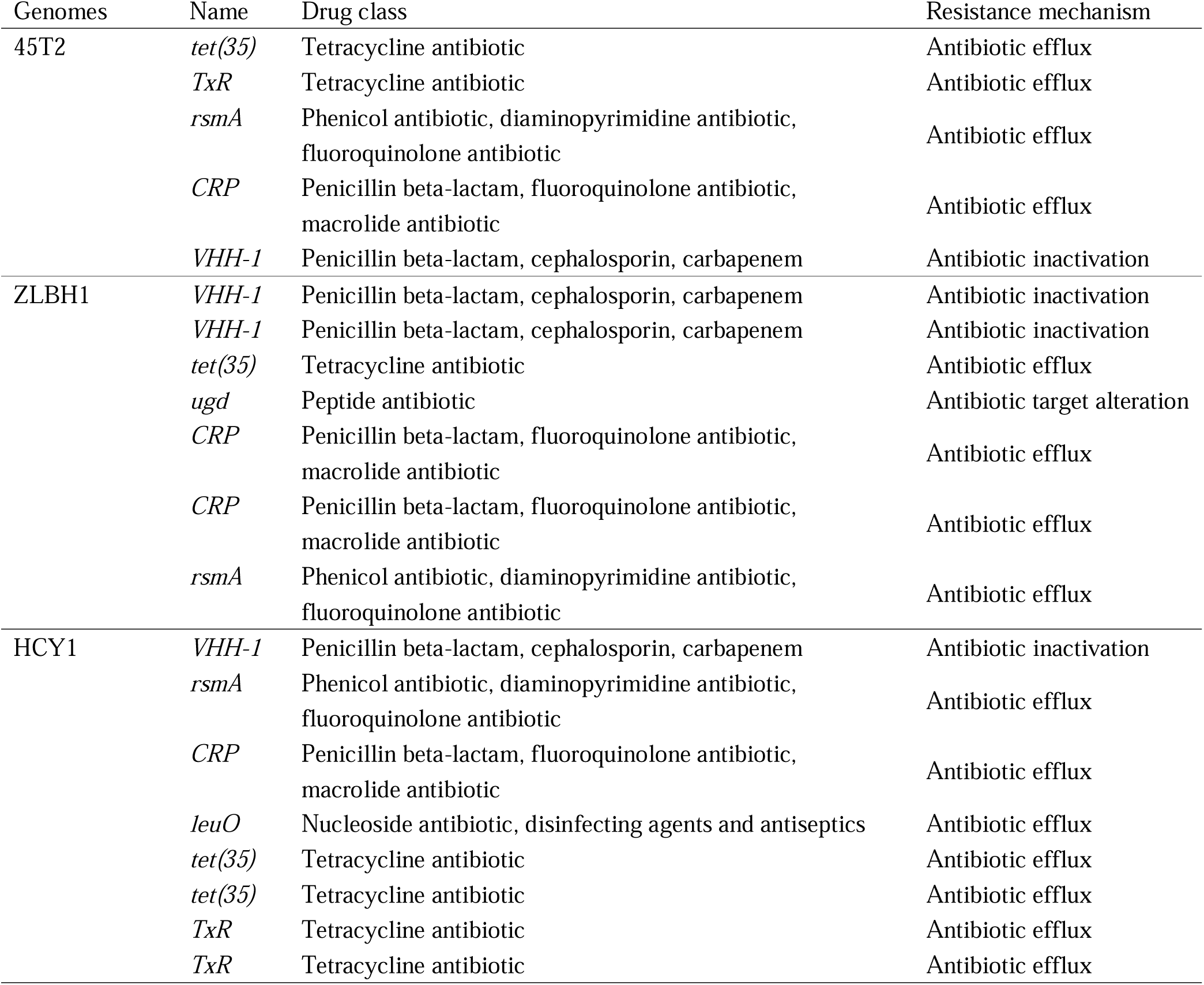
Antimicrobial resistance genes analysis of the *V. harveyi* strains 45T2, ZLBH1, and HCY1 in the CARD database with BLAST.

### 3.9 Virulence genes analysis

To further elucidate the differences in virulence genes among the three strains, we analyzed their virulence gene profiles (Fig. S1). Similarly, all virulence genes of HCY1 were located on the chromosomes, and HCY1 harbored more virulence genes than strains ZLBH1 and 45T2. Both highly virulent strains (HCY1 and ZLBH1) carried multiple siderophore-related genes (*entB*, *entF*, and *dhbE*), which likely contribute to their improved survival. Furthermore, both highly virulent strains possessed more complete type III secretion system (T3SS) and type VI secretion system (T6SS) gene sets. Although all three strains contained the major hemolysin gene *tlh* of *V. harveyi*, HCY1 uniquely carried the extracellular protease gene *hap/vvp* and the RTX toxin system gene *rtxB*, which may confer stronger infectivity and lethality.

### 3.10 The genes in the plasmids contribute to the enhancement of virulence

Since no virulence genes or AMR genes were identified on the plasmids, we performed further analysis of the genes carried by the plasmids. Following annotation, most of the genes on the plasmids were identified as hypothetical proteins (Table S2), with 69 genes successfully annotated (Table S3). These genes could be classified into five major categories (Table 4), among which 13 genes belonged to transposases, indicating that the plasmids carried by HCY1 possess a strong capacity for horizontal transfer. Additionally, many of the genes contribute to bacterial DNA maintenance and repair (e.g., *rep*, *dnaN*, and *dam*), thereby enhancing the strain’s stability and survival. Multiple genes facilitate host recognition, adhesion, and colonization, such as *mcpA*, *bdlA*, and *pal*. In addition, genes contributing to bacterial virulence and immune evasion were identified, such as *esiB*, which encodes a secreted immunoglobulin A-binding protein that impairs neutrophil activation (Pastorello et al., 2013).

**Table 4:** The genes carried by the plasmids of HCY1.

| Categories | Related genes |
| --- | --- |
| Mobile genetic elements | IS3, IS4, IS5, IS66, ISAs1, Tn3 family transposase |
| DNA maintenance and repair | *glcR*, *rep*, *repE*, *dinG*, *uvrD*, *xerC*, *dam*, *hns*, *ihfA/B*, *dnaN*, *ligA*, *tus*, *noc*, *hupB* |
| Metabolism and transport | *cynR*, *btuB*, *alaE*, *hpt*, *maa*, *cof*, *sppA*, *ycel*, *yceJ*, *rnk*, *coaD*, *orn* |
| Colonization and chemotaxis | *mcpA*, *bdIA*, *pal*, *tra*, *cph2* |
| Virulence and immune evasion | *esiB*, *hcpC*, *pleD*, *dosC*, *pdeC* |

## 4. Discussion

Chinese bahaba is a fish species endemic to China and is also listed as a Class I nationally protected wild animal. It holds exceptionally high value due to its highly valuable swim bladder, which is regarded as an important tonic in traditional Chinese medicine, especially in southern China (Sadovy de Mitcheson et al., 2019). Although significant breakthroughs have been made in the artificial breeding of Chinese bahaba, high mortality rates still occur during artificial culture. Therefore, understanding the major diseases affecting Chinese bahaba is crucial for protecting the captive population. In this study, the diseased fish exhibited pronounced skin ulcers, ascites, and gastroenteritis, and these clinical signs were highly consistent with those observed in typical vibriosis (Ina - Salwany et al., 2019). Through pathogen isolation, identification, and artificial challenge experiments, *V. harveyi* was confirmed as the causative agent of this disease outbreak. Given the special protected status of Chinese bahaba, we selected Miiuy croaker, its closest taxonomically related species, to conduct infection assays (Cui et al., 2024). Although this approach does not fully comply with Koch’s postulates, it is the most feasible strategy for studying diseases in endangered protected aquatic animals. A similar alternative strategy has been accepted in disease research on another critically endangered species, the Chinese sturgeon (*Acipenser sinensis*). Deng et al. (2022) used Dabry’s sturgeon (*Acipenser dabryanus*), the closest relative of the Chinese sturgeon, as a substitute host for challenge tests to investigate diseases caused by *Plesiomonas shigelloides* and *Citrobacter freundii*. Within 10 days post-infection, the miiuy croaker exhibited 100% cumulative mortality, accompanied by typical clinical signs including skin ulcers and ascites, which were consistent with those observed in naturally infected fish. These findings further confirmed the high virulence of this *V. harveyi* strain. To the best of our knowledge, this is the first systematic report of a disease outbreak in cultured Chinese bahaba, providing valuable insights for the disease control and population conservation of this species.

As an opportunistic pathogen, *V. harveyi* poses a significant threat to a variety of cultured aquatic animals (Zhang et al., 2020). It has been reported that *V. harveyi* infection in Pacific white shrimp (*Litopenaeus vannamei*) can result in mortality rates exceeding 80% (Soto-Rodriguez et al., 2012), causing substantial economic losses to the aquaculture industry. Additionally, *V. harveyi* infection caused pathological changes in multiple tissues and organs. In black rockfish (*Sebastes schlegeli*) infected with *V. harveyi,* presenting with skin ulcers, histopathological changes included hepatocyte vacuolation, glomerular atrophy, intestinal epithelial necrosis, and muscle fiber lysis (Gu et al., 2025). Infected hybrid grouper (*Epinephelus fuscoguttatus*♀×*E. lanceolatus*♂) also exhibited severe muscle necrosis and scale loss (Zhu et al., 2018). In barramundi (*Lates calcarifer*) infected with *V. harveyi*, head kidney necrosis, focal necrosis of peripancreatic cells accompanied by inflammatory cell infiltration, hepatocyte vacuolation, gill epithelial hyperplasia, and lamellar fusion were observed (Samsing et al., 2023). Schrama et al. (2025) observed that in broodstock Senegalese sole (*Solea senegalensis*) infected with *V. harveyi*, hemorrhagic lesions occurred in both the gills and skin, along with marked congestion and necrosis in the liver and spleen. Similar clinical signs (e.g., ulcer, gastroenteritis, and ascites) and histopathological changes (e.g., necrosis and inflammation in multiple tissues and organs) were also consistently observed in Chinese bahaba in this disease outbreak. These findings further corroborate the substantial threat posed by *V. harveyi* to aquaculture animals.

Since the discovery of antibiotics, they have been widely used in agriculture, animal husbandry, and aquaculture (Batuman et al., 2024; Done et al., 2015). The most effective method for treating bacterial diseases remains the use of antibiotics. In recent years, studies on the antimicrobial resistance profiles of *V. harveyi* isolates from aquaculture have revealed resistance to multiple antibiotics. Isolates from shrimp farms in India were found to be resistant to ampicillin, penicillin G, cefaclor, ciprofloxacin, ofloxacin, erythromycin, streptomycin, and vancomycin, while remaining susceptible to norfloxacin, chloramphenicol, and gentamicin (Stalin and Srinivasan, 2016). Three *V. harveyi* strains isolated from Pacific white shrimp farms in Malaysia all exhibited resistance to five antibiotics, including oxytetracycline, amoxicillin, ampicillin, colistin sulphate, and oleandomycin (Nurhafizah et al., 2021). The strain obtained from a seahorse (*Hippocampus kuda*) farm in China was resistant to multiple β-lactam antibiotics, such as ampicillin, penicillin, and cefazolin, as well as to polymyxin B and sulfisoxazole (Xie et al., 2020). The *V. harveyi* strain isolated from diseased Chinese bahaba in this study also exhibited resistance to multiple β-lactam and tetracycline antibiotics, as well as to enrofloxacin. However, the strain remained susceptible to florfenicol and trimethoprim-sulfamethoxazole, suggesting that these two antibiotics could be candidates for clinical treatment for this disease outbreak. Further genomic analysis revealed that HCY1 carried eight resistance genes, which primarily conferred resistance to β-lactams and tetracyclines, consistent with its phenotypic resistance profile. In addition, the strain harbored a unique resistance gene, *leuO*, a regulatory gene that may be associated with resistance to nucleoside antibiotics, disinfectants, and antiseptics. It is speculated that this may be the result of selective pressure from the long-term use of disinfectants in the aquaculture farm.

Bacterial infection of the host primarily relies on the virulence factors carried by the bacteria. Motility and chemotaxis are critical during the initial stages of infection (Yang and Defoirdt, 2015), as they determine whether the bacteria can successfully recognize the host. Adhesion is another essential step for successful infection (Cai and Arias, 2017; Finlay and Falkow, 1997), during which bacterial flagella also play an important role, such as the adhesion of *V. alginolyticus* to mucus (Chen et al., 2008). Besides, the bacteria colonize the host and form biofilms, which are also necessary for successful infection (Cai et al., 2013; Nurhafizah et al., 2021; Zhang et al., 2014). The type III secretion system (T3SS) is a needle-like structure commonly associated with the virulence of Gram-negative bacteria (Morot et al., 2025). It functions by injecting effector proteins synthesized within the bacterial cell directly into host cells, thereby causing cell lysis (Zhao et al., 2018). Studies have demonstrated that T3SS enabled *V. harveyi* to colonize and survive on the gills of European abalone (*Haliotis tuberculata*), and overproduction of the T3SS contributed to hemocyte death (Morot et al., 2025). Like the T3SS, the type VI secretion system (T6SS) is a protein delivery apparatus widely present in Gram-negative bacteria and is considered a key virulence system in *Vibrio* species (Boyer et al., 2009; Wan et al., 2024). In addition, pathogenic *V. harveyi* can produce various lytic enzymes, such as hemolysins and proteases, which damage host cells and release nutrients, thereby facilitating enhanced survival within the host. Meanwhile, siderophores are also considered critical virulence determinants in many *Vibrio* species (Darshanee Ruwandeepika et al., 2012). Comparative genomic analysis revealed that the *V. harveyi* strain isolated in this study carried many virulence-associated genes, including a more complete T3SS and T6SS, the hemolysin gene *tlh*, the extracellular protease gene *hap/vvp*, and the RTX toxin system gene *rtxB*. In addition, the strain harbored multiple siderophore-related genes (*entB*, *entF*, and *dhbE*). These virulence genes likely contribute to the strong pathogenicity of HCY1 and the high mortality observed in infected fish, which was further supported by the 100% mortality rate following artificial challenge.

Plasmids, as one of the most important mobile genetic elements (MGEs), facilitate horizontal gene transfer (Sobecky and Hazen, 2009). Plasmids not only enhance the genetic diversity of *V. harveyi* but also promote the rapid horizontal gene transfer of genes associated with virulence, antimicrobial resistance, and environmental stress responses (Deng et al., 2019). Studies have shown that some *V. harveyi* strains carry no plasmids, while others harbor multiple plasmids. For instance, the highly virulent strain N8T11 isolated from large yellow croaker carried five plasmids, which significantly contribute to its virulence (Wang et al., 2025). In this study, whole-genome sequencing and assembly of strain HCY1 revealed the presence of two plasmids of different sizes. Although no AMR genes or virulence genes from the CARD and VFDB databases were annotated on either plasmid, further analysis indicated that both plasmids still carry a substantial number of genes associated with enhancing virulence, including genes related to bacterial colonization, DNA repair, and immune evasion. For example, the methyl-accepting chemotaxis protein *mcpA*, a member of the methyl-accepting chemotaxis proteins (MCPs), upon binding to chemotactic ligands, generates chemotactic signals that are relayed through a series of chemotaxis proteins to the flagellar motor, enabling the bacterium to move toward favorable environments. This facilitates host seeking and colonization (Hida et al., 2015). Biofilm dispersion protein *bdlA* facilitates the transition of bacteria from a biofilm state to a single-cell planktonic state, and this phenotypic shift promotes host infection (Li et al., 2014). Peptidoglycan-associated lipoprotein *pal* constitutes a critical component of the Tol-Pal protein system (Godlewska et al., 2009). Low expression of the *pal* gene in *Escherichia coli* has been shown to attenuate virulence and enhance the survival rate of mice (Hellman et al., 2002). In addition, the secretory immunoglobulin A-binding protein *esiB* can inhibit neutrophil chemotaxis, which helps the bacterium evade immune clearance and reduces immune cell recruitment to the infection site, thereby enhancing bacterial survival within the host (Pastorello et al., 2013). We also identified multiple transposase systems within the plasmids, including the IS and Tn3 family transposases. These elements, through frequent transposition and homologous recombination, can significantly promote the horizontal gene transfer of virulence factors among strains, which may be a key factor in the evolution of this strain into a highly virulent strain.

## 5. Conclusion

In this study, we systematically reported for the first time a disease outbreak caused by *V. harveyi* infection in cultured Chinese bahaba through bacteriological, histopathological, and comparative genomic analyses. The results indicated that *V. harveyi* infection caused significant pathological damage in multiple tissues and organs of Chinese bahaba, particularly severe necrotizing gastroenteritis in the gastrointestinal tract. Additionally, strain HCY1 exhibited resistance to multiple antibiotics, including β-lactams and tetracyclines. Comparative genomic analysis revealed that HCY1 carried various AMR genes primarily conferring resistance to β-lactams and tetracyclines, which was highly consistent with the antimicrobial susceptibility testing results. Moreover, artificial infection with HCY1 resulted in a cumulative mortality of 100% within 10 days post-challenge. It was found that HCY1 possessed complete T3SS and T6SS gene clusters, along with virulence-associated genes such as *tlh* and *hap/vvp*. Furthermore, the plasmid harbored by HCY1 carried multiple genes that might enhance its virulence, such as *mcpA*, *pal*, and *esiB*, further corroborating its high virulence phenotype. In summary, our findings provide valuable insights for the disease control and population conservation of Chinese bahaba.

## Authors’ contributions

Liang Zhong: Writing-Original Draft, Methodology, Software, and Formal analysis. Ruoxi Zhu: Investigation and Conceptualization. Yuxuan Zhang: Software and Formal analysis. Lin Yan: Data curation and Resources. Song Sun: Investigation. Kuoqiu Yan: Conceptualization and Investigation. Wenlong Cai: Writing-Reviewing and Editing, Funding acquisition, Conceptualization, and Project administration.

## Acknowledgments

We thank Jingyuan Ruan in Guangdong Bluegen Marine Biotechnology Co., Ltd for assisting in the collection of diseased samples. We also acknowledge all authors for their contributions to this study.

