## Supplementary Table 1 and Supplementary Figure 1 for "Isolation, identification, and comparative genomic analysis of *Vibrio harveyi* as a causative agent for a skin ulcer disease of endangered fish Chinese bahaba (*Bahaba taipingensis*)"

**Supplementary materials**

Table S1: Sequencing and assembly statistics of strain HCY1.

| Strain | HCY1 |
| --- | --- |
| Total number of reads | 2,630,297 |
| Sum length (bp) | 1,563,393,716 |
| Average length (bp) | 594.4 |
| Max length (bp) | 303,284 |
| GC (%) | 44.37 |
| Coverage (×) | 254.02 |
| Genome size (bp) | 6154598 |
| Number of contigs | 4 |
| Number of plasmids | 2 |
| N50 | 3588647 |
| N90 | 2250360 |
| Number of CDSs | 9189 |
| Number of rRNA | 37 |
| Number of tRNA | 133 |

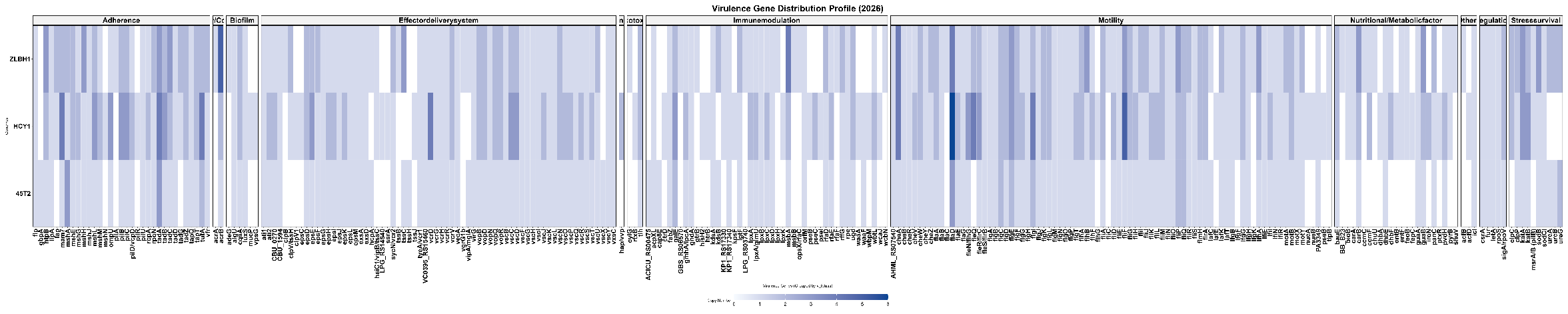

Figure S1: The virulence gene analysis of these three strains (ZLBH1, HCY1, and 45T2).
